# Temporal context actively shapes EEG signatures of time perception

**DOI:** 10.1101/2020.03.17.995704

**Authors:** Atser Damsma, Nadine Schlichting, Hedderik van Rijn

## Abstract

Our subjective perception of time is optimized to temporal regularities in the environment. This is illustrated by the central tendency effect: when estimating a range of intervals, short intervals are overestimated whereas long intervals are underestimated to reduce the overall estimation error. Most models of interval timing ascribe this effect to the weighting of the current interval with previous memory traces *after* the interval has been perceived. Alternatively, the *perception* of the duration could already be flexibly tuned to its temporal context. We investigated this hypothesis using an interval reproduction task in which human participants (both sexes) reproduced a shorter and longer interval range. As expected, reproductions were biased towards the subjective mean of each presented range. EEG analyses showed that temporal context indeed affected neural dynamics during the perception phase. Specifically, longer previous durations decreased CNV and P2 amplitude and increased beta power. In addition, multivariate pattern analysis showed that it is possible to decode context from the transient EEG signal quickly after both onset and offset of the perception phase. Together, these results suggest that temporal context creates dynamic expectations which actively affect the *perception* of duration.

**Significance Statement:** The subjective sense of duration does not arise in isolation, but is informed by previous experiences. This is demonstrated by abundant evidence showing that the production of duration estimates is biased towards previously experienced time intervals. However, it is yet unknown whether this temporal context actively affects perception or only asserts its influence in later, post-perceptual stages as proposed by most current formal models of this task. Using an interval reproduction task, we show that EEG signatures flexibly adapt to the temporal context during perceptual encoding. Furthermore, interval history can be decoded from the transient EEG signal even when the current duration was identical. Thus, our results demonstrate that context actively influences perception.

## Introduction

The way humans experience time is not only driven by the current stimulus, but is also influenced by previous experiences. According to Bayesian observer models, humans integrate noisy sensory representations (the likelihood) with previously learned stimulus statistics (the prior distribution). This is illustrated by the temporal context or central tendency effect: when presented with a range of intervals, short intervals are overestimated and long intervals are underestimated (Jazayeri & Shadlen, 2010). Furthermore, the prior distribution has been shown to be dynamically updated, such that more recent intervals have a greater influence on the current estimate (Dyjas, Bausenhart, & Ulrich, 2012; Taatgen & van Rijn, 2011; Wiener, Thompson, & Branch Coslett, 2014). Although there is abundant behavioral evidence for Bayesian integration in human time perception (Acerbi, Wolpert, & Vijayakumar, 2012; Cicchini, Arrighi, Cecchetti, Giusti, & Burr, 2012; Gu, Jurkowski, Lake, Malapani, & Meck, 2015; Hallez, Damsma, Rhodes, van Rijn, & Droit-Volet, 2019; Jazayeri & Shadlen, 2010; Maaß, Riemer, Wolbers, & van Rijn, 2019; Maaß, Schlichting, & van Rijn, 2019; Roach, McGraw, Whitaker, & Heron, 2017; Schlichting et al., 2018; Shi, Church, & Meck, 2013), its temporal locus and neural underpinnings are not yet understood.

Computational models of interval timing often (implicitly) assume that only after perception has completed, the noisy interval percept is weighted with previous memory traces representing the prior (e.g., Di Luca & Rhodes, 2016; Jazayeri & Shadlen, 2010; Taatgen & van Rijn, 2011). Alternatively, however, prior experience might actively affect perception, as evidenced by recent behavioral (Cicchini, Benedetto, & Burr, 2020; Cicchini, Mikellidou, & Burr, 2017; Zimmermann & Cicchini, 2020), fMRI (St. John-Saaltink, Kok, Lau, & De Lange, 2016) and single neuron findings (Sohn, Narain, Meirhaeghe, & Jazayeri, 2019). Specifically, Sohn et al. (2019) showed that neurons in the prefrontal cortex of monkeys exhibited different firing rate patterns based on the prior during interval estimation.

In humans, evidence is now emerging that electroencephalography (EEG) signatures in timing tasks are modulated by recently perceived durations. In a bisection task, longer prior durations led to a larger amplitude of the contingent negative variation (CNV) and increased beta oscillations power (Wiener, Parikh, Krakow, & Coslett, 2018; Wiener & Thompson, 2015). Crucially, however, these studies required an active comparison to the standard interval, in which EEG signatures have been shown to reflect an adjustment of the decision threshold (Ng, Tobin, & Penney, 2011; see also Boehm, Van Maanen, Forstmann, & van Rijn, 2014). Any context-based changes in these signatures might reflect updating of the comparison process. It is therefore still an open question what the temporal locus of the context effect is: Does the prior exert its influence in post-perceptual stages or are purely perceptual processes already affected by previous experiences?

We tested the influence of temporal context in an interval reproduction task, which allowed us to distill EEG signals during the perception phase in which no decision or motor response was required that could yield fallacious conclusions regarding the effect of context effects during perception. Participants reproduced two different interval ranges (the *short* and the *long context*). The ranges shared one interval (the *overlapping interval*), providing a condition in which the physical stimulus was the same, but the temporal context was different. We show that temporal context affects three EEG signatures that have previously been associated with time perception during the perception phase: the CNV and beta oscillations, but also the offset P2, which has been shown to predict subjective interval perception better than the CNV (Kononowicz & van Rijn, 2014; Kruijne, Olivers, & van Rijn, 2021). A data-driven approach reveals that temporal context can be decoded from transient neural dynamics during the perception phase using multivariate pattern analysis (MVPA). Together, these results show that temporal context actively shapes the perception of duration, falsifying most current formal theories of interval timing.

## Materials and Methods

### Participants

Twenty-seven healthy adults (22 females; age range 18 - 33 years, *M* = 21.33, *SD* = 3.78 years) participated in the experiment for course credits in the University of Groningen Psychology program or monetary compensation (€ 14). Two participants were excluded from analysis during pre-processing due to excessive artifacts in the EEG data. The study was approved by the Psychology Ethical Committee of the University of Groningen (17141-S-NE). Written informed consent was obtained before the experiment. After the experiment, the participants were debriefed about the aim of the study.

### Stimuli and apparatus

Stimuli were presented using the Psychophysics Toolbox 3.0.12 (Brainard, 1997; Kleiner et al., 2007) in Matlab 2014b. Intervals were presented as continuous 440 Hz sine wave tones. These auditory stimuli were presented on Sennheiser HD 280 Pro stereo headphones at a comfortable sound level. Visual stimuli were presented in the center of the screen in Helvetica size 25 in white on a dark grey background using a 27-inch Iiyama ProLite G27773HS monitor with a 1920×1080 resolution at 100 Hz. The index-finger trigger buttons of a gamepad (SideWinder Plug & Play Game Pad, Microsoft Corporation) were used to record responses.

### Procedure

Participants performed an auditory interval reproduction task (Figure 1A). Every trial started with a central fixation cross with a uniform random duration between 2 and 3 s. Then, an exclamation mark was presented for 0.7 s, after which the auditory interval was presented (the *perception phase*) while the exclamation mark remained on the screen. To signal the next phase, the exclamation mark was replaced by a question mark which was presented for 1.5 s. Next, the continuous tone was presented again, with the question mark remaining on the screen, which participants had to terminate by pressing a button (the *reproduction phase*). Participants were instructed to match the duration of this second tone to the duration of the first tone as accurately as possible.

**Figure 1.**
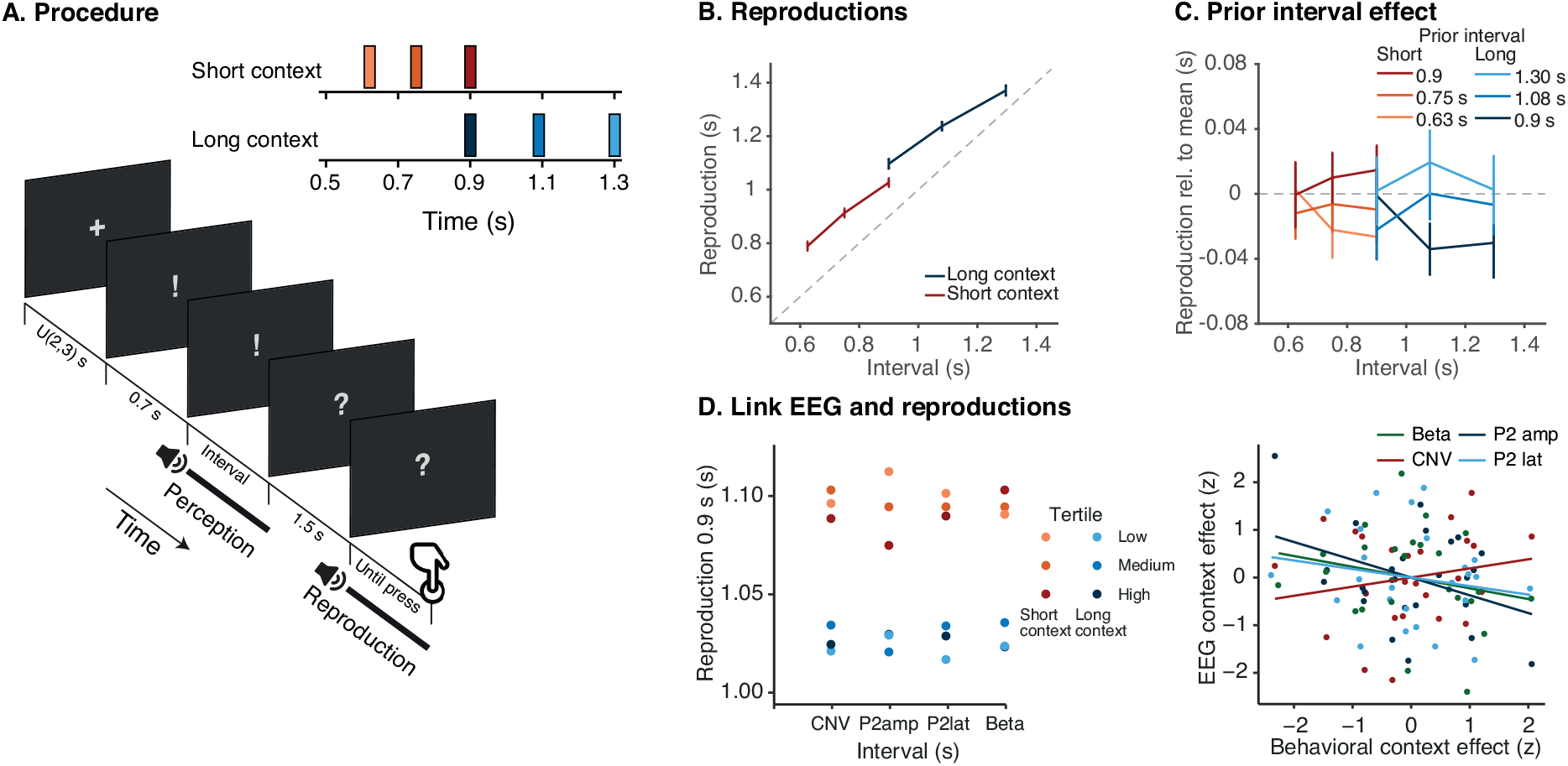
Task and behavioral results. A) Behavioral procedure of the experiment. Participants performed an interval reproduction task in which they heard a tone for a certain duration (*perception phase*). After an ISI of 1.5 s, they were asked to reproduce this duration by pressing a button to indicate the offset of the *reproduction phase*. In separate blocks, the perception phase consisted of three short or three relatively long durations (the *short* and the *long context*, respectively). One interval was presented in both contexts (the *overlapping interval* of 0.9 s). B) Average behavioral reproduction results. Error bars represent the standard error of the mean. C) Average reproduction of the overlapping interval (0.9 s) for the different intervals in the previous trial, relative to average reproduction in the context condition. Overall, reproductions were longer when the prior interval was longer. D) Link between the EEG signatures and reproductions. The left panel shows the reproduction of the overlapping interval for relatively low, medium, and high values (i.e., tertiles) of the CNV amplitude, P2 amplitude, P2 latency, and beta power. The right panel shows the correlation between participants’ behavioral context effect and their context effect in the different EEG signatures (all values were *z*-scored). Dots represent individual participants, while the lines represent linear regression lines.

The task involved two different interval ranges, the short context (0.625 s, 0.75 s, and 0.9 s) and the long context (0.9 s, 1.08 s, and 1.296 s) (Figure 1A). Crucially, there was an *overlapping interval* that was presented in both contexts (0.9 s). The experiment consisted of four blocks, two of which used intervals of the short context, and two of which used intervals of the long context. Block order was counterbalanced across participants, with the constraint that the context would alternate every block. Within a block, each duration of the short or the long context was presented 30 times, amounting to a total of 90 trials per block and 360 trials over the whole experiment. The presentation order was random, with the constraint that every possible subsequent pair of intervals was presented equally often (i.e., first-order counterbalancing). The hand needed for reproduction was switched after two blocks. Prior to each block, participants were instructed which hand (i.e., which gamepad button) they would use to terminate the duration and which set of intervals would be presented (termed set A and set B for the short and long context, respectively; see also Maaß, Schlichting, et al., 2019), while they were not informed about the relative durations or distributions associated with the sets (i.e., that the sets were associated with a short and long interval range). Participants could take self-timed breaks between blocks. Prior to the experiment, participants performed two practice trials with durations outside the range of both context conditions (0.4 s and 2 s). Experiment scripts are available at https://osf.io/sgbjz/.

### EEG acquisition

EEG signals were recorded from 62 Ag/AgCl electrodes, placed in accordance with the international 10-20 system (WaveGuard EEG cap, eemagine Medical Imaging Solutions GmbH, Berlin, Germany). The ground electrode was placed onto the left side of the collarbone and the mastoids served as location for the reference electrodes. The electrooculogram (EOG) was recorded from the outer sides of both eyes and from the top and bottom of the left eye. Data was collected at a sampling frequency of 512 Hz using a TMSi Refa 8-64 amplifier. Before the experiment, impedances of all electrodes were reduced to below 5kΩ. Participants were instructed to blink only between trials and not to move during the experiment.

### EEG pre-processing

EEG pre-processing was performed using the FieldTrip toolbox (Oostenveld, Fries, Maris, & Schoffelen, 2011). EEG data was re-referenced to the averaged mastoids and filtered using a Butterworth IIR band-pass filter with a high-pass frequency of 0.01 Hz and a low-pass frequency of 80 Hz. Subsequently, trial epochs were created from −1 s until 6 s relative to the onset of the perception phase. Artifacts were corrected using independent component analysis (ICA). Epochs that exceeded an amplitude range of 120 μV were removed from the dataset. On average, 10.72% (*SD* = 6.10) of the 360 trials were discarded.

### Data Analysis

#### Behavioral analysis

Reproductions lower than 0.1 s and higher than 2 s were excluded from analysis (0.2% of the data). To test whether reproductions were influenced by context, we fitted a linear mixed model (LMM) using the *lme4* package (Bates, Mächler, Bolker, & Walker, 2015) in R (R Core Team, 2016), including interval, context, their interaction and prior interval (i.e., the interval in the previous trial) as fixed factors. To facilitate interpretation of the results, interval and prior interval were centered at 0.9 s and the factor context was recoded using effect coding (−0.5 for short and 0.5 for long context). In addition to the random intercept of participant, we sequentially added random slope terms and tested whether they improved the model with a likelihood ratio test. We will here report the results of the best fitting model, which included random slopes for interval and prior interval.

#### ERP analysis

All EEG analyses reported here focused on the perception phase. An overview of the EEG results in the reproduction phase is available in the supplementary materials (section 1) at https://osf.io/sgbjz/. The CNV and beta signatures in the reproduction phase show trends that are qualitatively similar to the perception phase, although they appear to be less strong.

#### CNV

The CNV analysis was performed on a fronto-central electrode cluster (electrodes Cz, C1, C2, FCz, FC1, FC2) (Kononowicz & van Rijn, 2014; Ng et al., 2011). A 10 Hz Butterworth low-pass filter was applied and the ERP was baselined to the average signal in the 0.1 s window before interval onset. To test the effect of global context during the perception phase, we compared the ERP of the overlapping interval in the short and the long context using a cluster-based permutation test (Maris & Oostenveld, 2007) in the window 0-1.2 s from interval onset. The permutation test assessed whether the difference was different from zero by computing 100.000 permutations using the *t*-statistic, controlling for multiple comparisons with a cluster significance threshold of *p* < .05. To assess the influence of the prior interval on CNV, we calculated the average amplitude in the time window that showed CNV differences in the previously mentioned permutation test (0.3-1.01 s), per participant, context and prior interval for the overlapping interval. Next, we tested an LMM predicting this amplitude, including context and prior interval as fixed factors, and participant as a random intercept term.

#### Offset P2

The P2 analysis focused on the EEG signal averaged over the same fronto-central electrode cluster as the CNV analysis, to which a 1–20 Hz Butterworth band-pass filter was applied to minimize CNV-based contamination (cf., Kononowicz & van Rijn, 2014). The ERP was baselined to the average signal in the 0.1 s window around interval offset (cf., Kononowicz & van Rijn, 2014). Similar to the CNV analysis, the ERPs of the overlapping interval in the short and the long context were compared using a cluster-based permutation test in the window 0-0.5 s after interval offset. Next, we calculated P2 amplitude was as the average amplitude between 0.14 and 0.3 s after interval offset (this window was based on Kononowicz & van Rijn, 2014). We fitted an LMM predicting P2 amplitude, with interval, prior interval, and context as fixed factors, and participant as a random intercept term. The random slope of interval improved the fit and was added to the model. P2 latency was calculated as the 50% area latency - the time point at which half of the area under the curve is reached - within the same window (Liesefeld, 2018; Luck, 2005). P2 latency was analyzed using an LMM with the same fixed factors as the P2 amplitude model.

Because the 1 Hz high-pass filter might induce artifactual effects of opposite polarity before the actual peak (Tanner, Morgan-Short, & Luck, 2015), we also performed the P2 analysis on the data without additional filtering (that is, besides the band-pass filter between 0.01 Hz and 80 Hz applied during pre-processing). We found similar qualitative results, which are reported in the supplementary materials section 2.1 at https://osf.io/sgbjz/.

#### Multivariate pattern analysis

To investigate transient neural dynamics in more detail, we tested whether it is possible to decode global and local context through MVPA of the EEG signal. Following Wolff, Kandemir, Stokes, and Akyürek (2020), we used a sliding window approach in which the EEG fluctuations were pooled over electrodes and time. A window of 50 data points (98 ms) was moved across the signal in steps of 8 ms, separately for each channel. Within the window, the signal was down-sampled to 10 samples (by taking the average over 5 samples) and baseline-corrected by subtracting the mean within the window from all 10 individual samples.

To decode whether an overlapping-interval trial was presented in the short or the long context, the 10 samples per electrode in each time window served as input for 5-fold cross-validation. In each fold, we calculated the Mahalanobis distance (De Maesschalck, Jouan-Rimbaud, & Massart, 2000; Wolff, Jochim, Akyürek, & Stokes, 2017; Wolff et al., 2020) between the test trials and the averaged signal of the short and long context, using the covariance matrix estimated from the training trials with a shrinkage estimator (Ledoit & Wolf, 2004). To make the distance estimates more reliable, the 5-fold cross-validation was repeated 50 times and results were averaged. The eventual decoding distances were smoothed with a Gaussian smoothing kernel (*SD* = 16 ms). To test whether the distance between conditions was significantly different from zero, a cluster-based permutation test was performed.

A similar analysis was performed to decode the duration of the prior interval from the neural dynamics in the current trial. For the overlapping interval, the Mahalanobis distance between every test trial and the average of the prior interval conditions was calculated. This resulted in six difference time series for each condition (including the 0.9 s condition for each context separately and the difference with the trial’s own condition). In this way, we aimed to determine whether the distance was higher when the difference between the prior interval condition of the test trial and the other possible prior interval conditions was larger. Next, for every time point, we performed a linear regression on the Mahalanobis distance, using the absolute difference between prior interval conditions (in seconds) and the difference between context (coded as 0 or 1) as predictors, allowing us to disentangle the effect of sequential and global context on transient neural dynamics. A cluster-based permutation test was performed on the resulting slope values for prior interval and context, to test whether they deviated from zero (using a one-sided *t*-test).

To investigate which electrodes are most informative in decoding the context of an overlapping interval trial, we performed channel-wise decoding: the procedure to decode global context outlined above was performed separately for every electrode. Topographies were created to show the average decoding accuracy at the different electrodes during time windows in which the Mahalanobis distance resulting from the context decoding procedure outlined above (i.e., using all electrodes) was significantly higher than zero.

Because the context conditions were blocked in our experimental design, the decoding accuracy might have been inflated by nonstationarities in the EEG signals, which lead to stochastic dependence between trials (Lemm, Blankertz, Dickhaus, & Müller, 2011). Post-hoc, we controlled for this notion by calculating the Mahalanobis distance between the different blocks, for each participant. This allowed us to differentiate between the distances between blocks that were presented in the first and second half of the experiment, and thereby, to test whether the original decoding results could be due to within-block similarities beyond context. In this way, we compared the Mahalanobis distance between the trials in a particular block and the ‘same context’ and ‘different context’ block in the other half of the experiment. We found that the results were qualitatively similar to the original analysis, with significant differences between the short and long context immediately after interval onset and after interval offset (analysis details and full results can be found in the supplementary materials section 2.2. at https://osf.io/sgbjz/).

#### Time-frequency analysis

To assess oscillatory power during the perception phase we performed a time-frequency analysis using a single Hanning taper with an adaptive time window of 6 cycles per frequency in steps of 15 ms for frequencies from 4 to 40 Hz, with the amount of spectral smoothing set to 1. We calculated the absolute power from the baseline window of −0.2-0 s relative to interval onset. The analysis was again focused on fronto-central electrodes (Cz, C1, C2, FCz, FC1, FC2). Similar to the CNV analysis, all time-frequency analyses were performed on the overlapping interval to isolate the effect of context while keeping the actual stimulus constant.

Per participant, for every time-frequency point, we fitted a linear regression model including prior interval (a continuous variable ranging from the shortest to the longest interval in seconds) and context (short and long context coded as 0 and 1, respectively) as predictors (following an approach similar to Wiener et al., 2018). For every time-frequency point, this resulted in two slope values, expressing the relative influence of the global context and the previous interval. Next, a one-sample *t*-test against zero was performed for the two slope values at each time-frequency point, which was corrected for multiple comparisons using cluster-based permutation (Maris & Oostenveld, 2007). The statistical testing was performed on the frequency range of 8-30 Hz to include alpha power (8–14 Hz; Kononowicz & van Rijn, 2015) and beta power (15–30 Hz; e.g., Haegens et al., 2011; Jenkinson & Brown, 2011; Kononowicz & van Rijn, 2015) during the time window of 0-1.2 s after interval onset.

#### Linking EEG signatures and behavior

We tested in two ways whether EEG signatures during the perception phase predicted behavioral reproductions. First, we computed *single trial* values of CNV amplitude, P2 amplitude, P2 latency and beta power. Following the methods described above, for every trial, CNV amplitude was calculated as the average EEG signal in the window 0.3-1.01 s after interval onset, P2 amplitude as the average between 0.14 and 0.3 s after interval offset, P2 latency as the 50% area latency in the same window, and beta power was calculated as the average power in the time window 0.48-0.84 s after interval onset and the frequency range 23-30 Hz, which was based on the permutation test. CNV, P2 amplitude, P2 latency, and beta power values that deviated more than 4 SD from the average were excluded from analysis (0.06%, 0.01%, 0.00% and 0.46% of the trials, respectively). Similar to the behavioral analysis described above, reproductions shorter than 0.1 s and longer than 2 s were also excluded from analysis. Next, we computed four LMMs with reproduction as the dependent factor, and CNV amplitude, P2 amplitude, P2 latency, and beta power as fixed factors, respectively. Similar to the analyses described above, the CNV and beta power analyses were focused on the overlapping interval trials. To control for the effect of context on both EEG signatures and behavior, context and prior interval were also added as fixed factors to the models. The P2 analysis included all intervals, so here, interval was entered as an additional fixed factor. In all models, participant was included as a random intercept term, and adding random slopes did not improve the model fit in any of the models.

Second, in addition to the single trial analysis, we looked at individual differences: Do participants who show a large context effect in the EEG signatures also show a large behavioral context effect? To this end, for the overlapping interval, we estimated the behavioral context effect (i.e., the difference in reproduction between the long and short context) for each participant, and compared it to the context effect of CNV amplitude, P2 amplitude and latency, and beta power, quantified as described in the previous paragraphs. To assess whether these measures were related for each participant, we performed a one-tailed Pearson’s correlation test between the individual behavioral context effects and the EEG context effects.

## Results

### Behavioral results

Figure 1B shows the average reproductions for the different intervals. The results of the LMM showed that the reproductions increased with duration (*β* = 0.77, *SE* = 0.03, *t* = 24.33, *p* < .001). We found a significant effect of global context, showing that reproductions were longer in the long compared to the short context (*β* = 0.05, *SE* = 0.01, *t* = 7.23, *p* < .001). In addition, the increase with duration (i.e., the slope) was lower for the long compared to the short context (*β* = −0.18, *SE* = 0.03, *t* = −7.01, *p* < .001). Besides the global context effect, reproductions were longer when the interval in the previous trial was longer (*β* = 0.08, *SE* = 0.02, *t* = 3.41, *p* = .002). Figure 1C shows the reproductions for the different previous intervals, relative to the average reproduction.

### ERPs

#### CNV

Figure 2A and 2B show the average fronto-central ERP during the perception phase for the different intervals in the short and the long context, respectively. In addition, Figure 2D shows a direct comparison between the short and the long context of this ERP for the overlapping interval (0.9 s). The cluster-based permutation test showed that the CNV was more negative in the short than in the long context in the time windows 0.30-0.65 s (*p* = .004) and 0.71-1.01 s (*p* = .003). Thus, while the actual interval was the same, CNV amplitude during perception differed depending on the temporal context.

**Figure 2.**
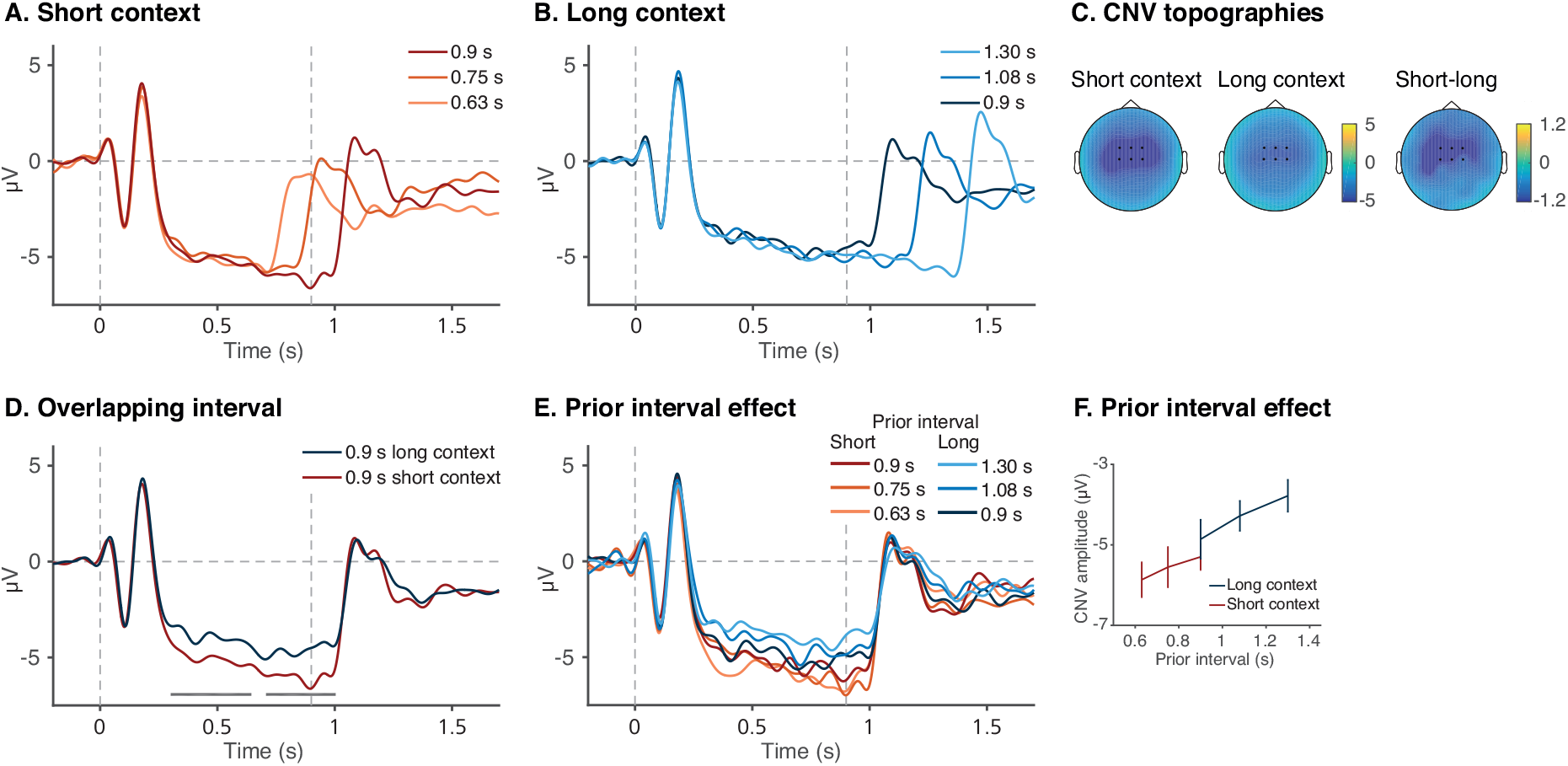
Average ERPs at the fronto-central cluster (Cz, C1, C2, FCz, FC1, FC2) relative to the onset of the perception phase for the different durations in the short (A) and long (B) context. In all panels, vertical grey dashed lines indicate interval onset and offset of the overlapping interval (0.9 s). C) Topographies of the overlapping interval (0.9 s), for the short context, long context, and their difference, during the window of significant difference as indicated by the cluster-based permutation test. D) Average ERP of the overlapping interval (0.9 s) in the short and the long context. Grey horizontal bars indicate significant differences according to the cluster-based permutation test. E) Average ERP of the overlapping interval, split up according to the interval in the previous trial. Red and blue lines show whether the overlapping interval appeared in the short or the long context, respectively. F) Average CNV amplitude for the middle interval, in the time window of significant difference between the short and long context, for the different previous intervals. Error bars represent the standard error of the mean.

Figure 2E shows the average ERP for the overlapping interval, split for the different previous durations, and Figure 2F shows the average CNV in the 0.3-1.01 s window for the different prior interval conditions. The LMM results showed that CNV amplitude at the overlapping interval became less negative with longer previous trials (*β* = 2.50, *SE* = 0.97, *t* = 2.57, *p* = .011). There was no evidence for an additional significant effect of context (*β* = 0.43, *SE* = 0.42, *t* = 1.03, *p* = .308), suggesting that the global context effect on CNV might be largely driven by the previous trial. Post-hoc, we tested whether including the interaction between context and prior interval improved the model fit, but a likelihood ratio test showed that this was not the case (*χ*^2^(1) = 0.10, *p* = .750).

### Offset P2

#### Amplitude

Figure 3A and 3B shows the offset P2 for the different intervals in the short and the long context, respectively. Figure 3D directly compares the P2 for the overlapping interval in the short and long context. The cluster-based permutation test showed that the amplitude was higher for the short compared to the long context in the window 0.11-0.3 s. Figure 3F shows the average P2 amplitude as a function of interval and context. The LMM showed that the P2 increased with duration (*β* = 5.56, *SE* = 0.62, *t* = 8.97,*p* < .001), but that the intercept was significantly lower for the long compared to the short context (*β* = −0.87, *SE* = 0.28, *t* = −3.10, *p* = .002). Figure 3E and 3G (left panel) show the effect of the prior interval on P2 amplitude for the overlapping interval. In line with the global context effect, the model showed that the P2 decreased with longer previous intervals (*β* = −1.71, *SE* = 0.51, *t* = −3.35, *p* = .001). Together, these results show that P2 amplitude reflects the actual duration, as well as the global and local context in which the duration appeared.

**Figure 3.**
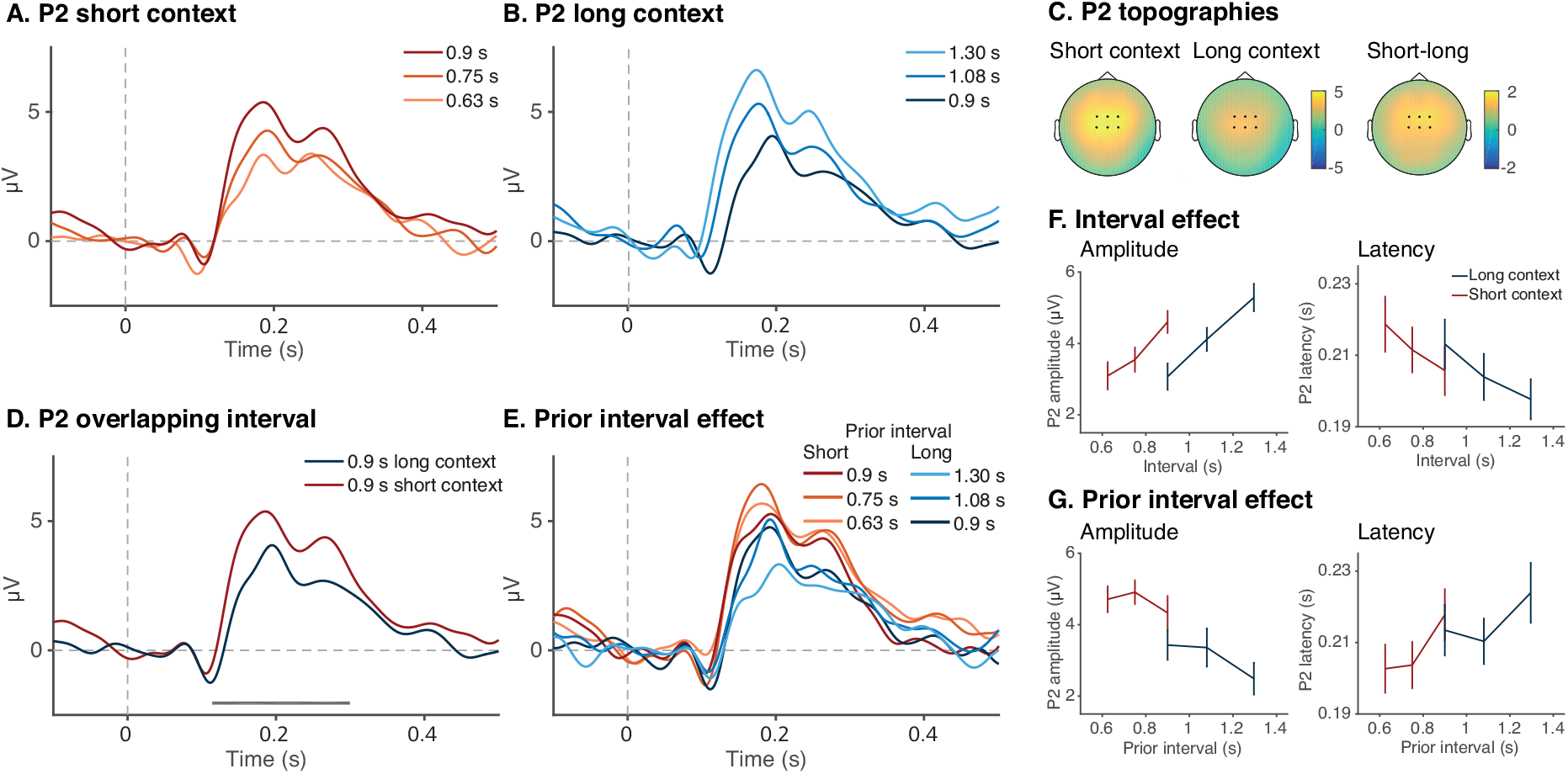
Amplitude and latency of the P2 at the fronto-central cluster (Cz, C1, C2, FCz, FC1, FC2) after the offset of the perception phase. A, B) Grand average ERPs baselined at the offset of the perception phase in the short and the long context, respectively. C) Topographies of P2 amplitude of the overlapping interval (0.9 s), for the short context, long context, and their difference, during the window of significant difference as indicated by the cluster-based permutation test. D) Average ERP of the overlapping interval (0.9 s) in the short and the long context. Grey horizontal bars indicate significant differences according to the cluster-based permutation test. F) Effect of interval on P2 amplitude and latency. The left panel shows P2 amplitude, calculated as the average amplitude in the window 0.14-0.3 s after interval offset for every participant and interval. The right panel shows P2 latency, calculated as the 50% area latency in the same window. G) Effect of the prior interval on P2 amplitude (left) and latency (right). In all figures, error bars represent the standard error of the mean.

#### Latency

Figure 3F (right panel) shows that P2 latency decreased with the duration of the current interval, which was confirmed by the LMM predicting latency (*β* = −0.04, *SE* = 0.01, *t* = −3.66, *p* < .001). There was no evidence that P2 latency was affected by the context, as the fixed factors contex and prior interval did not reach significance (*ps* > .247). In summary, whereas P2 amplitude reflects the current duration and the general and sequential temporal context, P2 latency only decreases with longer current durations.

### Multivariate pattern analysis

Figure 4A shows the decoding accuracy for the overlapping interval. The permutation test showed a positive cluster immediately after interval onset (0-0.17 s; *p* = .009) and after interval offset (0.99-1.37 s; *p* < .001). Figure 4C shows the topographies of the channel-wise decoding results during these two clusters, which reflects high parietal and left-lateralized decoding accuracy and high fronto-central and right-lateralized decoding accuracy, respectively. Figure 4B shows the slope value of prior interval in the regression analysis predicting Mahalanobis distance. The permutation test showed that there was no evidence for significant clusters for the slope of prior interval or context in the regression analysis (*p* = .999), showing that MVPA could not distinguish between prior interval conditions based on the transient EEG signal.

**Figure 4.**
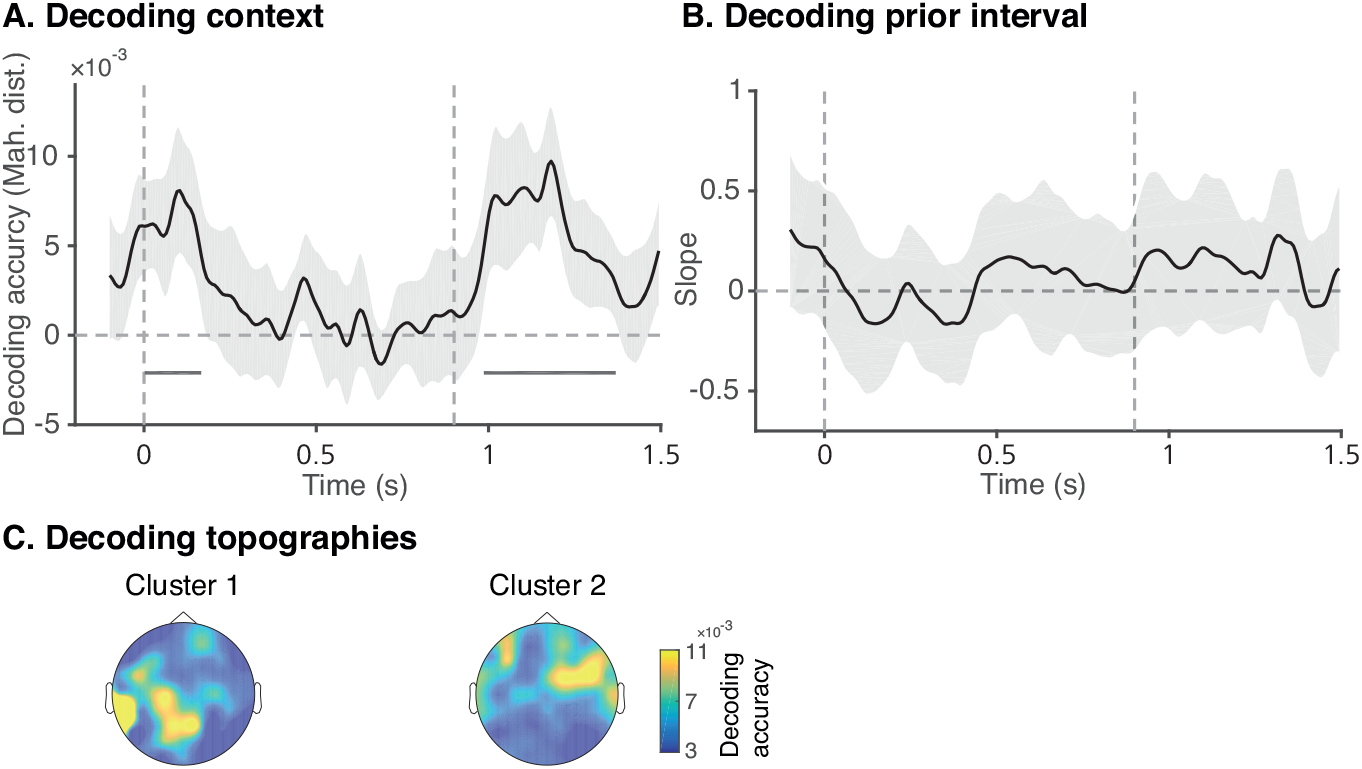
Decoding accuracy relative to the onset of the perception phase. A) Decoding accuracy of context for the overlapping interval as represented by the Mahalanobis distance. Grey horizontal bars indicate a significant difference from zero according to the cluster-based permutation test. Error shading represent 95% CI of the mean. B) Decoding accuracy of prior interval in the overlapping interval, represented by the slope value of the regression of Mahalanobis distance with prior interval and context as predictors. C) Topographies of channel-wise context decoding accuracy for the overlapping interval, during the first significant cluster in panel A (left) and the second cluster (right). Colors represent the decoding accuracy in Mahalanobis distance.

### Time-frequency analysis

To assess oscillatory power during the perception phase, we calculated a linear regression of frequency power at fronto-central electrodes with context (short vs long) and prior interval as predictors for every time-frequency point during the overlapping interval. Figure 5A shows the slope values representing the effect of context on the power of the different frequencies over time. We found a positive cluster in the window 0.48-0.84 s after interval onset in the 23-30 Hz frequency range (*p* = .045), indicating increased beta power in the long context compared to the short context (see the outlined area in Figure 5A). This beta effect is further illustrated in Figure 3C, which shows the average power in the 23-30 Hz over time, for the overlapping interval in the short and long context. Figure 5B shows the slope values for prior interval, for which the permutation test indicated no evidence for a cluster of slopes different from zero (*ps* > .051). In summary, these results suggest that fronto-central beta power was higher in the long compared to the short context, while there was no evidence for a similar influence of the previous trial.

**Figure 5.**
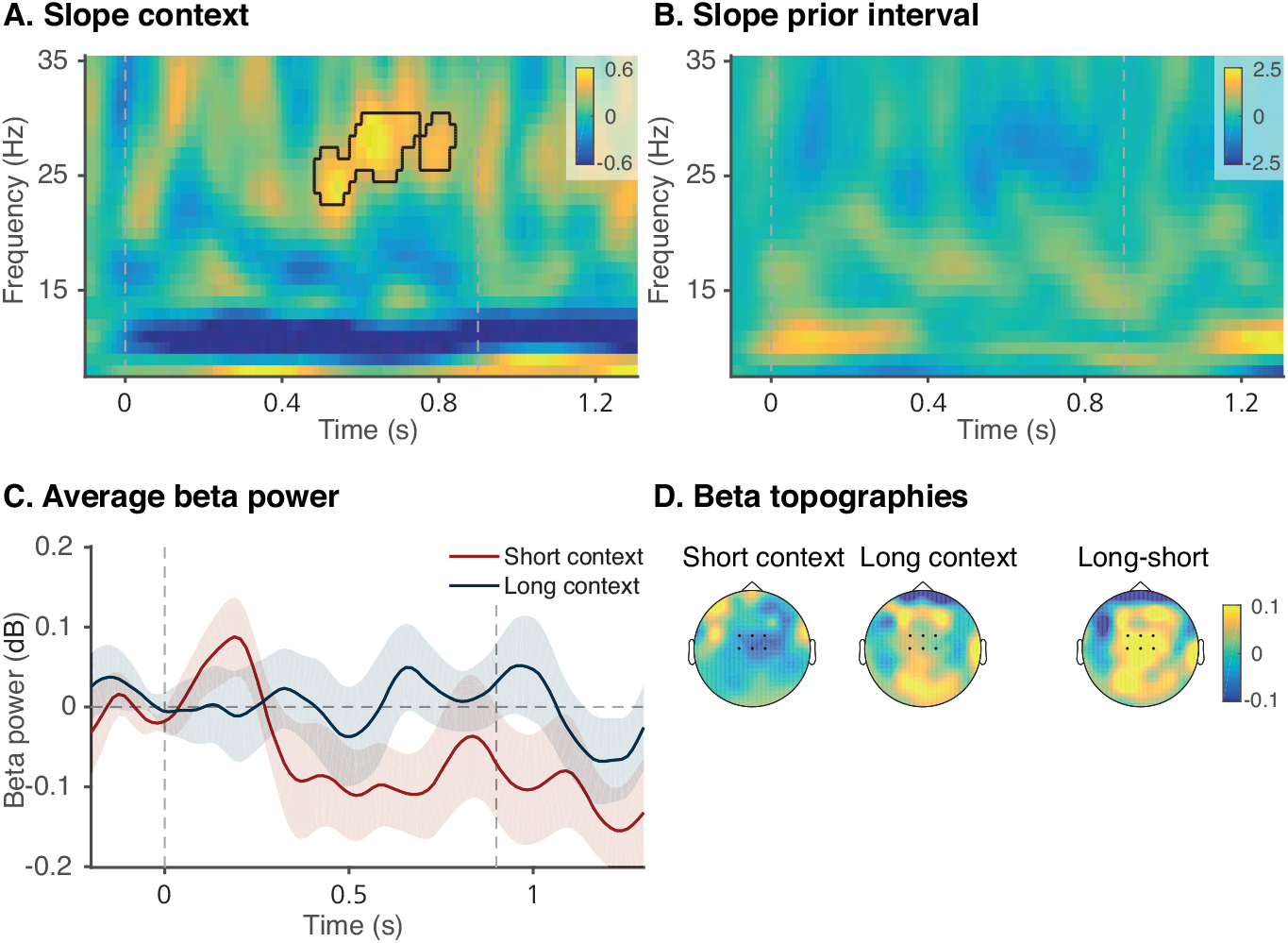
Slope values of regression on frequency power at fronto-central electrode cluster (electrodes Cz, C1, C2, FCz, FC1, FC2) relative to the onset of the perception phase. A) Slope values of the factor Context (short vs long) in the regression analysis at every time-frequency point. The outlined area marks a significant cluster according to the cluster-based permutation test performed in the time window 0-1.2 s and the frequency window 8-30 Hz. B) Slope values of the factor prior interval in the regression analysis predicting power. There was no evidence for significant clusters. In both panels, vertical dashed grey lines indicate the onset and offset of the perception phase. C) Average beta power in the time and frequency range of the significant cluster (23-30 Hz) for the short and long context for the overlapping interval. Error shading represents represent the standard error of the mean. D) Topographies of beta power for the overlapping interval, in the time and frequency range of the significant cluster, for the short context, the long context, and their difference.

### Linking EEG signatures and behavior

Figure 1C shows the effect of single-trial EEG signatures on reproductions of the overlapping interval. For illustration purposes, the single-trial EEG amplitudes and latencies were divided into tertiles (low/short, medium, high/long) for each participant and context, and the average reproduction was plotted for each tertile. The LMMs showed no evidence that single-trial CNV and beta power in the perception phase predicted reproductions in the reproduction phase (*β* = 0.0003, *SE* = 0.0004, *t* = 0.85, *p* = .395 and *β* = 0.0003, *SE* = 0.0019, *t* = −0.51, *p* = .614, respectively). This was also the case for P2 latency, with a trend towards shorter reproductions for later P2 peaks (*β* = −0.08, *SE* = 0.04, *t* = −1.80, *p* = .072). However, P2 amplitude after perception phase offset was predictive of that trial’s reproduction (*β* = −0.0010, *SE* = 0.0003, *t* = −3.45, *p* < .001). As the *β*-value indicates, higher P2 peaks were followed by shorter reproductions. Given that context, interval and prior interval were also included as fixed factors in the LMM, these results cannot be attributed to a mediating influence of context, and therefore suggest that trial-by-trial variation in P2 amplitude might be a reliable predictor of reproductions.

We additionally tested whether participants with a large behavioral context effect for the overlapping interval also showed a large context effect in the EEG signatures. This between-participant relationship between these measures is depicted in Figure 1D. Analysis showed that the individual behavioral context effect was correlated with the P2 amplitude difference between contexts (*r*(23) = −.37, *p* = .033). We found no evidence for a similar relationship with P2 latency (*r*(23) = −.18, *p* = .196), CNV amplitude (*r*(23) = .19, *p* = .180) or beta power (*r*(23) = −.22, *p* = .861). Thus, in line with the single trial analysis, P2 amplitude differences predict reproduction outcomes.

## Discussion

As the temporal locus of Bayesian computations in human time estimation is still unknown, we investigated whether temporal context actively influences neural signatures during the perception of time intervals. Behaviorally, we found that reproductions were biased towards the global temporal context as well as the duration in the previous trial. EEG results showed that CNV, P2 and beta power were modulated by previously perceived intervals, and that context could be decoded from transient brain dynamics at an early stage during perception. These results indicate that previously perceived durations actively affect EEG signatures during interval estimation, showing that prior experiences act directly on perception. This observation goes against the (implicit) assumption of time perception models that the likelihood is weighted with the prior only after perception. It is, however, in line with recent behavioral evidence showing that context asserts its influence at early sensory stages (Cicchini et al., 2020; Zimmermann & Cicchini, 2020). Our findings suggest that experiences with the global and recent temporal context actively calibrate cortical dynamics, in which the CNV and beta power may reflect the anticipation of stimulus duration, and the P2 component the active evaluation of the interval in the current context. Crucially, by focusing on the perception phase in a reproduction paradigm, this is the first work demonstrating context effects that are not linked to explicit motor preparation or response decisions.

Our findings argue against the idea that the CNV reflects the neural counterpart of the absolute accumulator in pacemaker-accumulator models (Casini & Vidal, 2011; Macar & Besson, 1985; Macar & Vidal, 2004; Macar, Vidal, & Casini, 1999; Macar & Vitton, 1982; Pfeuty, Ragot, & Pouthas, 2005), since no differences based on prior experience would be expected during the perception of an interval. Instead, we found that the CNV during the perception of the overlapping interval was more negative for the short compared to the long context, and for shorter previous durations. This is consistent with anticipation and preparation accounts of the CNV (e.g., Boehm et al., 2014; Elbert, 1993; Leuthold, Sommer, & Ulrich, 2004; Mento, 2013; Ng et al., 2011; Scheibe, Schubert, Sommer, & Heekeren, 2009) and pacemaker-accumulator models that propose adaptive spike rate accumulation (Simen, Balci, deSouza, Cohen, & Holmes, 2011): When interval offset is expected quickly after onset, CNV amplitude increases more rapidly. This adaptation is in line with studies showing a faster CNV development for relatively short foreperiods (Miniussi, Wilding, Coull, & Nobre, 1999; Müller-Gethmann, Ulrich, & Rinkenauer, 2003; Trillenberg, Verleger, Wascher, Wauschkuhn, & Wessel, 2000; Van der Lubbe, Los, Jaśkowski, & Verleger, 2004), shorter standard durations in an interval comparison task (Pfeuty et al., 2005), and after adaptation to a shorter interval (Li, Chen, Xiao, Liu, & Huang, 2017). The contextual adjustment of the speed with which the CNV develops suggests that neural populations in the supplementary motor area (SMA), which is typically associated with the CNV (e.g., Coull, Vidal, & Burle, 2016), can perform flexible temporal scaling based on the temporal context (Remington, Egger, Narain, Wang, & Jazayeri, 2018; Remington, Narain, Hosseini, & Jazayeri, 2018; Sohn et al., 2019), even in the absence of explicit motor preparation. The prior might calibrate the speed of neural dynamics through different initial states at the onset of the perception phase (Remington, Egger, et al., 2018; Sohn et al., 2019), as our multivariate pattern analysis showed that global context can be decoded from EEG dynamics immediately after the onset of the perception phase. Although the precise onset of significant decoding should be interpreted with caution since the moving window approach and low-pass filtering could smear out the accuracy over time (Grootswagers, Wardle, & Carlson, 2017), these results suggest that temporal context affects the instantaneous neural response to to-be-timed stimuli.

The active anticipation based on context was also indexed by the P2 component. Specifically, P2 amplitude increased with longer current durations, suggesting that it reflects hazard-based expectancy: the probability that the interval offset will occur, given that is has not yet occurred (Nobre, Correa, & Coull, 2007). This in line with previous studies showing that longer ISIs increase P2 amplitude (e.g., Pereira et al., 2014; Röder et al., 2000). Importantly, however, P2 amplitude decreased with longer previous durations, showing that the expectations are updated to the current temporal context, even on a trial-by-trial basis. These results complement previous studies showing that temporal expectancy modulates ERP amplitude (e.g., Kononowicz & van Rijn, 2014; Li et al., 2017; Todorovic & de Lange, 2012; Todorovic, van Ede, Maris, & de Lange, 2011; Wacongne et al., 2011). Interestingly, P2 amplitude at perception phase offset predicted interval reproductions, and participants’ behavioral context effect correlated with their context-based P2 effect. The lack of an equivalent CNV-effect highlights the predictive quality of the P2 (Kononowicz & van Rijn, 2014; Kruijne et al., 2021), and indicates that the neural state at the end of the perception phase sets the speed of cortical dynamics during reproduction (Sohn et al., 2019). Global context additionally influenced beta power, such that beta power was higher in the long compared to the short context, in line with effects of beta power in single trial analyses (Kononowicz & van Rijn, 2015). Although beta power has been proposed to reflect motor inhibition (Alegre et al., 2004; Hwang, Ghuman, Manoach, Jones, & Luna, 2014; Kononowicz & van Rijn, 2015; Kühn et al., 2004), and most studies on the link between beta power and timing have a strong motor component, our results suggest that synchronized beta oscillations also play a role during interval perception after which no immediate motor response is required. This finding complements recent studies showing that the accuracy and precision of time estimates depend on beta (Wiener et al., 2018) and alpha-beta coupling (Kononowicz, Sander, van Rijn, & Van Wassenhove, 2020). Additionally, the current global context effect on beta is in line with Wiener et al.’s finding that longer previous durations increased beta power in the current trial. It has to be noted, however, that we found no evidence for similar sequential effects on beta.

Besides the auditory stimuli which participants had to time, the current paradigm also consisted of visual stimuli that indicated the phase of the trial (i.e., perception or reproduction). The general overestimation we found in the behavioral results might potentially be explained by the integration of these visual stimuli in temporal estimation (Shi & Burr, 2016). Future studies might look further into potential modality differences in contextual calibration and their neural underpinnings (Rhodes, Seth, & Roseboom, 2018; Roach et al., 2017; Zimmermann & Cicchini, 2020). Furthermore, we found no significant decoding corresponding to the windows of CNV differences. This can be explained by the specific decoding method we employed, which focused on transient dynamics, filtering out the stable CNV activity by baselining within a moving window. In addition, decoding might be especially sensitive to stimulus onset and offset, with accuracy peaking shortly afterwards and slowly dropping as the neural synchronization declines (e.g., Wolff et al., 2017, 2020).

A comparison to Wiener and Thompson (2015), who found a larger CNV amplitude for *longer* prior durations, suggests that contextual ERP effects might be dependent on the specific experimental paradigm. In contrast to our reproduction experiment, their bisection task requires an active decision during perception, and the CNV has been shown to reflect this decision process by deflecting or plateauing after the standard interval in memory has been reached (Macar & Vidal, 2004; Ng et al., 2011; Pfeuty, Ragot, & Pouthas, 2003). A similar explanation could account for the different nature of our offset P2 effect compared to Kononowicz and van Rijn (2014), who found a V-shaped P2 amplitude attenuation in a temporal comparison task (but see Kruijne et al., 2021). This pattern reflects active comparison to the standard interval, which is not applicable to the current reproduction paradigm. In addition, the P2 measured in the current study shows similarities to the positive offset peak named the late positive component of timing (LPCt) (Gontier et al., 2009; Paul et al., 2011; Wiener & Thompson, 2015), although it has been argued that the P2 reflects perceptual predictive processes while the LPCt indexes decision making (Kononowicz, van Rijn, & Meck, 2016). The extent to which these components indeed reflect similar processes is still an open question, and their occurrence seems to depend on the specific nature of the task. Future studies might directly compare these neural differences in paradigms involving decision, motor or only perceptual timing requirements.

In conclusion, our results show that previous durations actively influence flexible neural dynamics during temporal encoding. These findings indicate that previous experiences in memory create expectations that in turn calibrate our perception of the environment. The adaptive influence of prior knowledge on perception could represent a more general Bayesian mechanism of magnitude estimation (Petzschner, Glasauer, & Stephan, 2015), falsifying a class of models that assume discrete, post-perceptual stages in which previous experiences exert their influence.

## Acknowledgements

This research has been partially supported by the EU Horizon 2020 FET Proactive grant TIMESTORM - Mind and Time: Investigation of the Temporal Traits of Human-Machine Convergence (Grant number 641100) and by the research programme Interval Timing in the Real World financed by the Netherlands Organisation for Scientific Research (NWO, Grant number 453-16-005, awarded to Hedderik van Rijn). The funding agencies had no involvement in the design of the study, the analysis of the data, writing of the report, or in the decision to submit the article for publication. We thank Sarah Maaß for fruitful discussions and programming the experiment, Ronja Eike for her help in data collection and preprocessing, Emil Uffelmann for his help in data collection, and Michael Wolff and Güven Kandemir for sharing their decoding knowledge with us.

